# Evaluation of *TH*-Cre knock-in cell lines for detection and specific targeting of stem cell-derived dopaminergic neurons

**DOI:** 10.1101/2020.07.10.193870

**Authors:** A Fiorenzano, J Nelander Wahlestedt, M Parmar

## Abstract

The focal and progressive degeneration of dopaminergic (DA) neurons in ventral midbrain has made Parkinson’s disease (PD) a particularly interesting target of cell-based therapies. However, ethical issues and limited tissue availability have so far hindered the widespread use of human fetal tissue in cell-replacement therapy. DA neurons derived from human pluripotent stem cells (hPSCs) offer unprecedented opportunities to access a renewable source of cells suitable for PD therapeutic applications. To better understand the functional properties of stem-cell derived DA neurons, we generated targeted hPSC lines with the gene coding for Cre recombinase knocked into the *TH* locus. When combined with flexed GFP, they serve as reporter cell lines able to identify and isolate TH+ neurons *in vitro* and after transplantation *in vivo*. These *TH*-Cre lines provide a valuable genetic tool to manipulate DA neurons useful for the design of more precise DA differentiation protocols and the study of these cells after transplantation in pre-clinical animal models of PD.

## Introduction

The degeneration of dopaminergic (DA) neurons in ventral midbrain (VM) is known to play a key role in the development of Parkinson's disease (PD). Attempts to design a cell therapy for PD based on replacing cells lost to the disease with new healthy DA neurons have spanned more than three decades_1_. Early trials using fetal VM tissue provided proof-of-concept that such an approach can be effective in slowing disease progression. However, the limited availability of fetal tissue as well as ethical and practical complications associated with its use have hampered the advancement of this therapeutic strategy and pointed to the need to identify alternative cell sources_2_. Human pluripotent stem cells (hPSCs) are widely expected to provide an alternative donor cell population available in unlimited quantities, and global efforts to use hPSC-derived DA neurons in PD patients are ongoing_3, 4_.

Previous experimental studies showed the function and innervation of stem cell-derived DA neurons in pre-clinical models of PD_5–9_. To further refine such studies and to track DA neurons during *in vitro* differentiation, we generated targeted cell lines with the gene coding for Cre recombinase knocked into the *TH* locus. When combined with flexed GFP, they serve as reporter cell lines allowing for the identification and isolation of TH neurons *in vitro* and after transplantation *in vivo*. These *TH*-Cre lines provide a robust and flexible genetic tool to manipulate DA neurons such as by inserting designer receptors exclusively activated by drugs (DREADDs) or channelrhodopsins for the specific activation or inhibition of DA neurons, or by performing rabies-based tracing from transplanted DA neurons. The ability to detect and specifically target DA neurons will represent a crucial step forward in the continuous efforts to design more precise DA differentiation protocols and to study these cells after transplantation in pre-clinical animal models of PD.

## Results

### Generation and validation of CRISPR/Cas9-mediated *TH*-Cre knock-in reporter cell lines

To knock Cre recombinase into the *TH* locus, we designed two different targeting approaches. One was based on knocking Cre into exon 1 of the *TH* gene (Fig. 1a), which has three isoforms (−001, −002, and −003) sharing the same start codon in exon 1. The other mimicked a previously reported *TH*-Cre rat model_10_, with the additional insertion of an internal ribosomal entry site (IRES)-Cre cassette in exon 14_11_ (Fig. 1b). Targeting of the donor templates was achieved using specific guide RNAs and CRISPR/Cas9 technology. We verified the correct insertion at both the 5’ and 3’ ends by PCR genotyping (Supplementary Fig. 1a,d). We used primer pairs binding outside TH donor vector homology arms and a second primer binding within sequences not contained in wild-type cells, thereby distinguishing targeted clones from those with random genomic insertions (Supplementary Fig. 1a,d and Supplementary Table1). DNA sequencing of PCR products confirmed precise insertion of the reporter gene without introducing errors, indicating that genomic editing was efficiently driven in *TH* locus (Supplementary Fig. 1c,f). To determine whether the insertion was present in one or both alleles, PCR genotyping was used to distinguish the wild-type from the modified allele and we found that *TH*-Cre and IRES-Cre knock-in clones were heterozygous and homozygous for the insertion, respectively (Supplementary Fig. 1b,e and Supplementary Table1). Based on these findings, we identified and selected two correctly targeted clones from the *TH*-Cre (A2 and D3) and the *TH*-IRES-Cre (B2 and C4) design for further analysis (Supplementary Fig. 1a,b,d,f). To assess if genome modification of the targeted *TH* locus impacted the ability of hPSCs to differentiate, we induced these four clones to DA progenitors following a well-established protocol_12_, _13_. The ventral and caudal midbrain patterning of these *TH*-Cre and *TH*-IRES-Cre targeted clones was compared to that of the parental cell line differentiated in parallel. We confirmed correct VM patterning by performing qPCR for key genes (*SHH, CORIN, FOXA2, OTX1, LMXA1, LMX1B, OTX2,* and *EN1*) (Fig. 1c and Supplementary Table2) and by assessing co-expression of LMX1a/FOXA2 and LMX1A/OTX2 at protein level (Fig. 1d). Taken together, these findings indicate that knock-in of Cre in *TH* locus does not alter the developmental potential of targeted cells to differentiate.

**Fig. 1.**
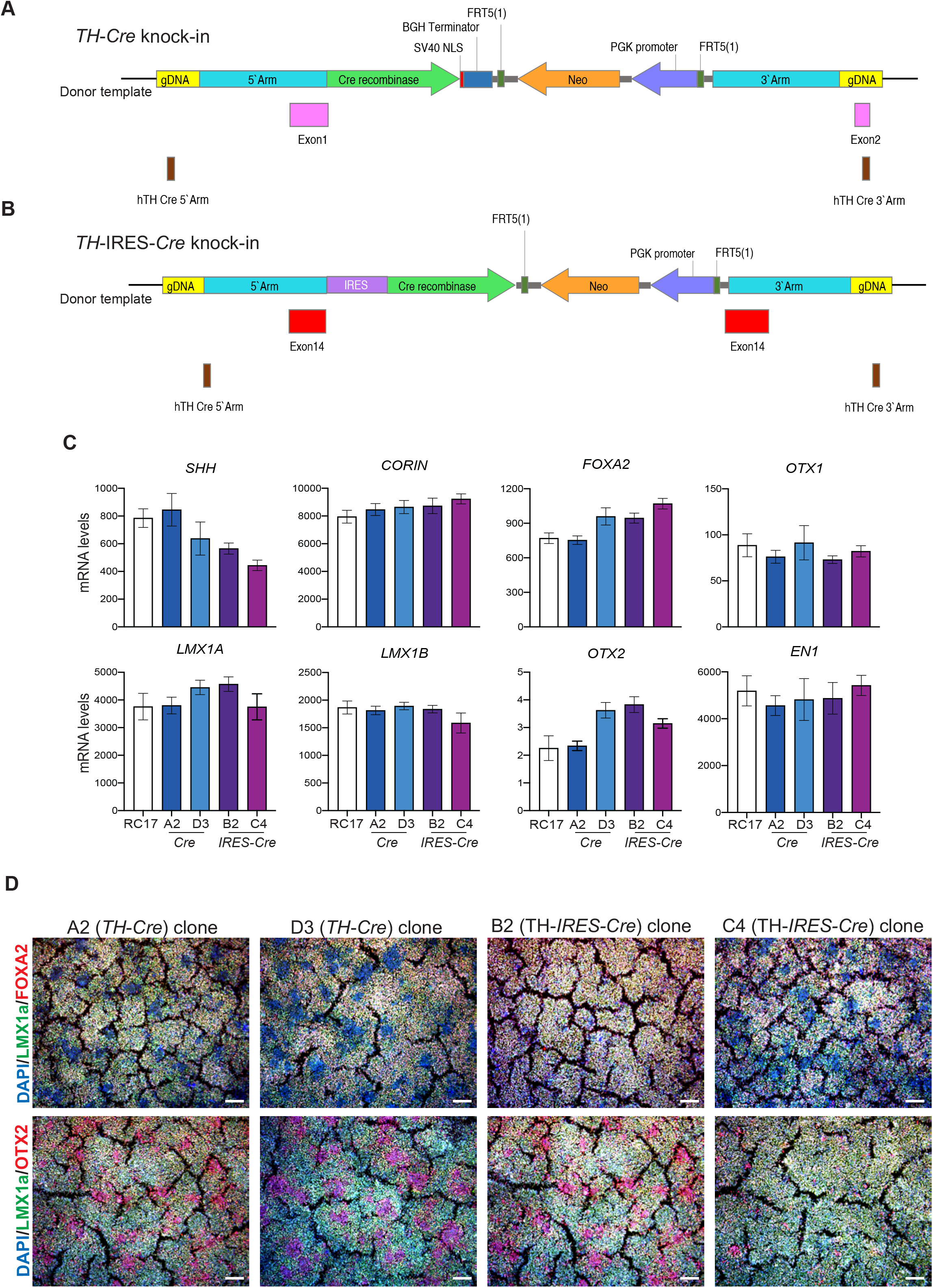
**a,** Schematic overview of targeting strategies to knock Cre into exon 1 **b,** or IRES-Cre into 3’UTR of human *TH* gene. **c,** qRT-PCR of selected markers at day 16 of cells patterned toward ventral midbrain (VM). Results are given as fold change over undifferentiated hPSCs. Data represent mean ± SEM of 3 independent experiments. **d,** Immunohistochemistry of LMX1a/FOXA2 and LMX1a/OTX2 at day 16 in Cre-targeted hPSC clones (A2 and D3), and of IRES-Cre targeted hPSC clones (B2 and C4) patterned toward VM. Scale bars, 100 μM. Nuclei were stained with DAPI.

### GFP expression reflects TH protein levels in *TH*-Cre knock-in reporter lines

To test the ability of selected transgenic hPSC-reporter cell lines to detect progenitor and mature DA neurons in live cultures, we transduced the cells with a Cre-inducible (DIO, or double-floxed inverted orientation) GFP expression cassette in undifferentiated hPSCs (Supplementary Fig. 2a). The specificity of flip-excision (FLEx)-GFP to TH+-expressing cells and the off-target function of this vector was first tested at protein level in undifferentiated hPSC culture.

FACS and immunofluorescence analysis detected no ectopic expression of GFP in absence of Cre (Suppl. Fig. 2) when transgenic lines were transduced with FLEx-GFP virus alone Supplementary Fig. 2b,d). In contrast, when hPSCs were co-transduced with both Cre and FLEx-GFP lentivirus, GFP was only detected in Cre-expressing cells (Supplementary Fig. 2c,e). We then analyzed the expression of TH, Cre, and GFP at day 28 after initiation of DA differentiation (Fig. 2a). Immunofluoresence confirmed that Cre expression mirrored TH protein levels, indicating that the targeting did not affect overall TH synthesis and stability (Fig. 2b,c). Subsequently, we found that GFP completely overlapped TH+ cells, confirming that TH and GFP are specifically co-expressed in both clones from the *TH*-Cre knock-in (Fig. 2b-c). By contrast, in the IRES-Cre line, GFP was found expressed both earlier and more broadly than TH in both independent B2 and C4 clones (Fig. 2d,e). We therefore focused the remainder of our analysis on the *TH*-Cre knock-in clones. We further differentiated these selected hPSC reporter lines into DA progenitors and used FACS to enrich TH+ cells in culture (Fig. 2f). Based on GFP expression, we sorted and collected GFP+ and GFP− cells, and then analyzed TH expression after 12 hrs. Immunofluorescence analysis confirmed that most TH neurons were recovered from the GFP+ cell population (Fig. 2g, h).

**Fig. 2.**
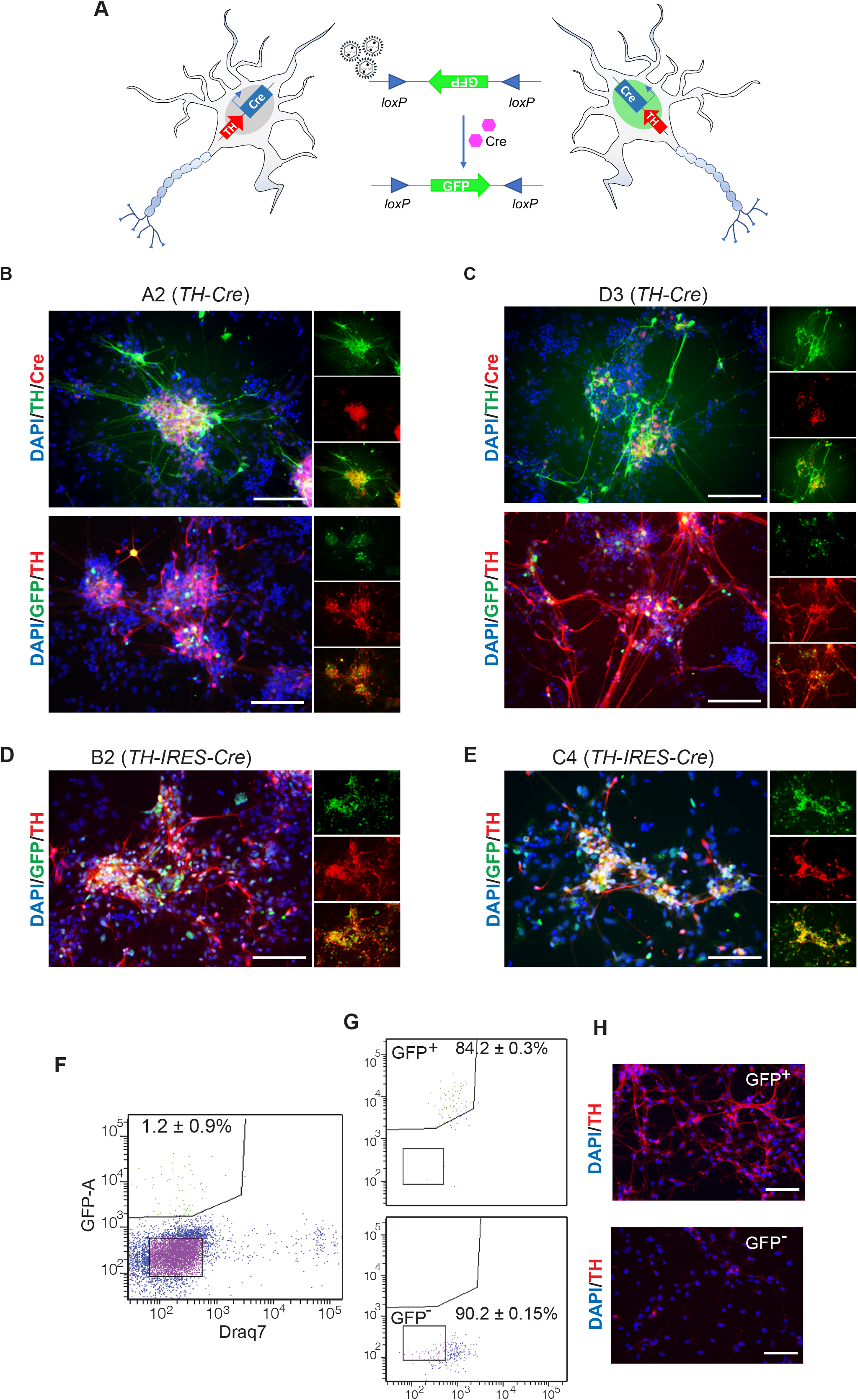
**a,** Schematic representation of FLEx-GFP in *TH*-Cre reporter cell line. **b,** Immunohistochemistry of TH/CRE and GFP/TH in A2 and **c,** in D3 *TH*-Cre targeted hPSC clones at day 28 during VM differentiation. Scale bars, 250 μM. Nuclei were stained with DAPI. **d,** FACS-based separation and **e,** post-sorting analysis of VM progenitor GFP+ and GFP− cells. Data represent mean ± SEM (*n*= 3). **f,** Immunohistochemistry of TH in GFP+ and GFP− FACS-separated cells.

### *TH*-Cre reporter cell line allows the study of developing and mature DA neurons

To determine whether mature and late DA neurons can be detected using the *TH*-Cre reporter cell line, we tested the persistence of GFP expression in terminally differentiated DA neurons cells by performing long-term *in vitro* and *in vivo* experiments (Fig. 3a,d). We transduced *TH*-Cre cells with LV-FLEx-ncGFP at DA progenitor state (day 12) and subsequently differentiated them *in vitro* until day 50 (Fig. 3a). Gene expression analysis for *TH*, *NURR1*, *PITX3*, *AADC*, *MAP2*, and *FOXA2* (Fig. 3b and Supplementary Table2) revealed proper DA differentiation patterning. In line with this finding, immunohistochemistry analysis confirmed the overlap between GFP and TH expression (Fig. 3c), demonstrating that targeting does not affect TH protein levels and that GFP is restricted to the TH expressing neurons. Using a similar strategy but with a nuclear GFP reporter (Fig. 3d), we transplanted day 16 progenitors into the striatum of 6-OHDA lesioned nude rats and then analyzed the cells 3 months after grafting by immunohistochemistry (Fig. 3e). This analysis corroborated that ncGFP is maintained also in mature TH neurons after grafting.

**Fig. 3.**
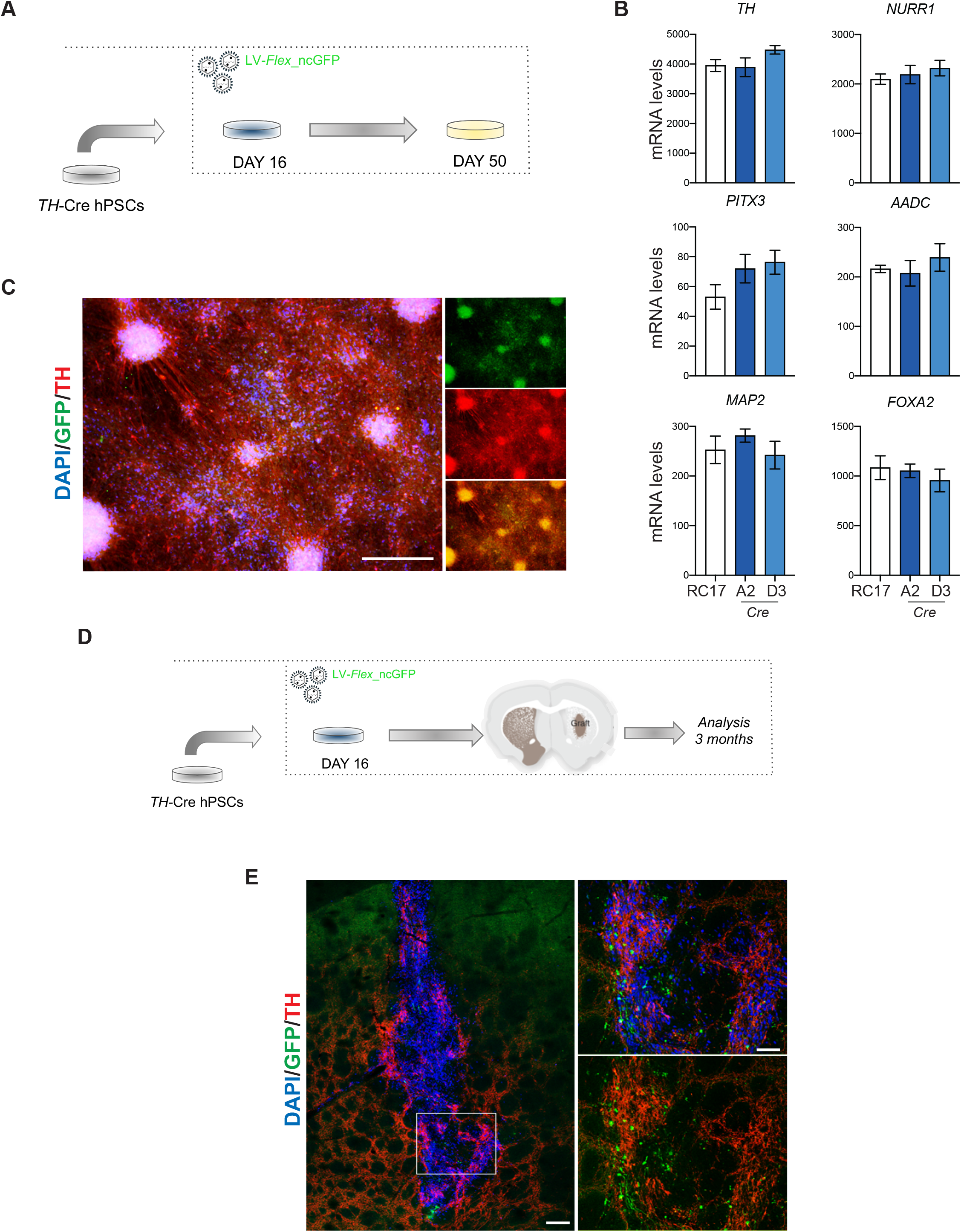
**a,** Schematic overview of *in vitro* experimental plan. **b,** Representative qRT-PCR of selected markers at day 50 of terminally differentiated VM cells. Results are given as fold change over undifferentiated hPSCs. Data represent mean ± SEM of 3 independent experiments. **c,** Immunohistochemistry of TH/GFP of A2 *TH*-Cre targeted hPSC clone at day 50 during VM differentiation. Scale bars, 250 μM. Nuclei were stained with DAPI. **d,** Schematic overview of in *vivo* experimental plan. **e,** Immunohistochemistry of TH/GFP in the graft core of VM-derived hPSC intrastriatal grafts 3 months post-transplantation. Insets show high-power magnifications of GFP+ DA neurons in the graft. Scale bars, 200 μM. Nuclei were counterstained with HUNU.

## Discussion

Here we describe the establishment of hPSC reporter lines where Cre is knocked into exon 1 of the *TH* gene. Using viral vectors where GFP and ncGFP (DOI) are transduced into pluripotent and/or progenitor cells, we showed that this design results in the faithful expression of GFP exclusively in TH+-expressing neurons *in vitro* and after transplantation *in vivo*. In this *TH*-Cre reporter line, targeting disrupts the *TH* gene and TH is only expressed from one allele. We also tested an alternative approach based on a *TH*-Cre rat model_10_ with the additional insertion of an IRES-Cre cassette in 3’UTR. This transgenic reporter line has the advantage that the endogenous *TH* gene is not disrupted and that a homozygous Cre knock-in is possible if high Cre levels are required. Although this design works well in rats _7, 10_, we found some leakage of expression outside *TH*+ neurons in stem cell cultures.

Knocking Cre into the gene locus instead of following the more conventional strategy of inserting a specific fluorescent reporter_14_ presents several advantages. *Firstly*, it allows for greater flexibility in the choice of reporter. Here we describe the use of GFP, which is useful when studying DA neurons *in vitro* or assessing morphology and projections *in vivo*, and of ncGFP, which can also be used for live-cell imaging, facilitates quantification, and allows for cell/nuclei sorting after transplantation has been performed with ubiquitously expressed GFP_15_. For other applications, any type of reporter could be inserted as required. Live reporters in DA neurons can also be used in more advanced techniques such as scRNAseq and Patch-seq analysis in order to combine assessment of genome-wide expression with neuronal activity and morphological characterization, thus facilitating the functional classification and mapping of DA neuronal subtypes. *Secondly*, many types of construct could be incorporated and used to refine *in vivo* studies where specifically targeting and studying the DA component of grafts is challenging_5, 6, 16_. *Thirdly*, the *TH*-Cre line can be adopted to selectively regulate DA neuron function in a bimodal manner, as previously performed with transplanted rat cells_7_.

We envision that this *TH*-Cre reporter cell line will serve as a valuable tool to refine stem cell differentiations and to better understand the functions and properties of DA neurons after transplantation.

## Supplementary Figure Legends

**S1.**
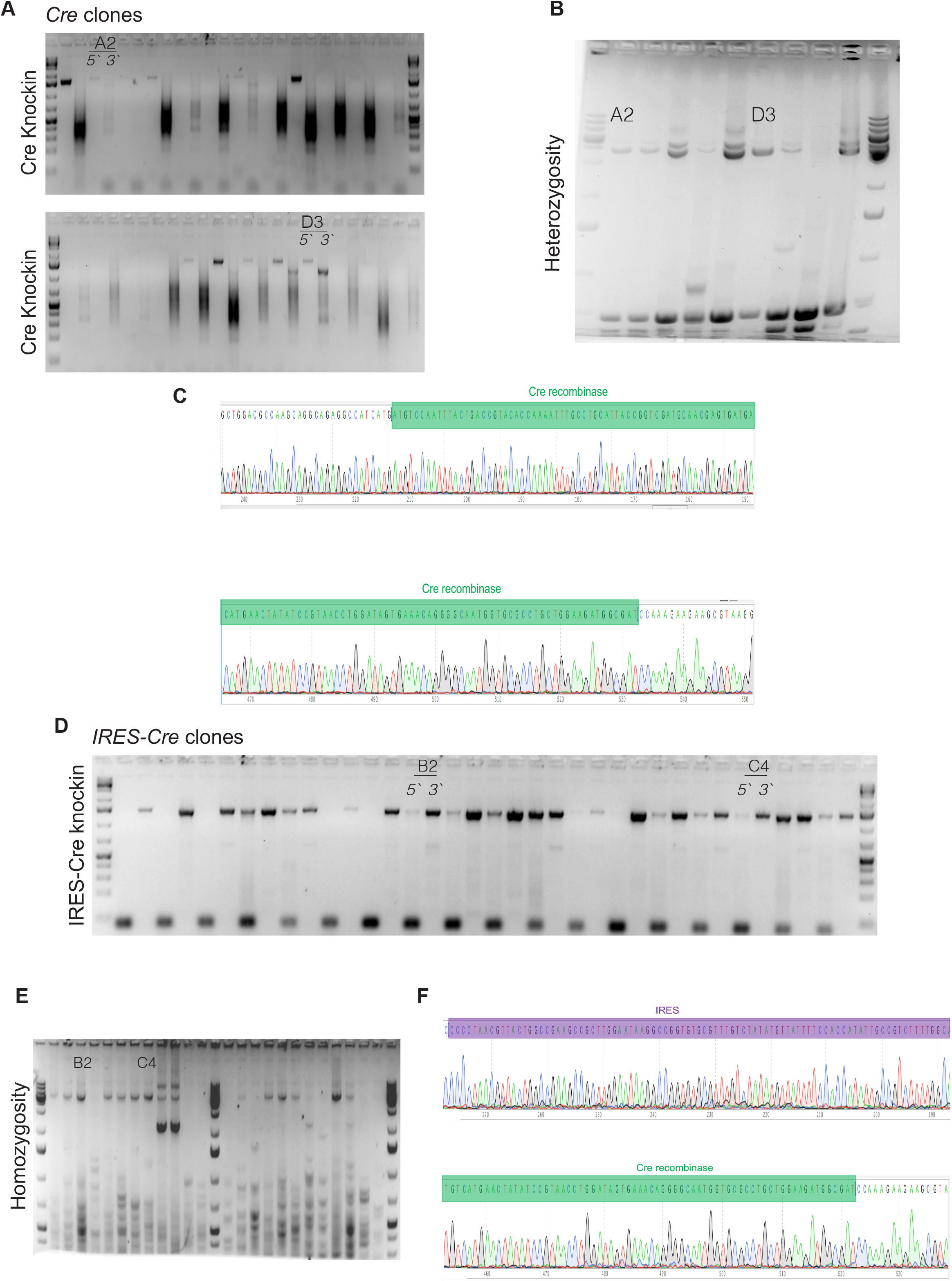
**a,** Genotyping PCR for 5′ arm (hTH3 UTR cassette 1R/Cre, 1522 bp) and 3′ arm (hTH3 UTR 3arm 2F/PGK, 1222 bp) of *TH*-Cre knock-in hPSC clones (A2 and D3). **b,** Genotyping PCR showing heterozygosity of targeted allele of *TH*-Cre hPSC clones (A2 and D3; knock-in 3.6 kb and wild type 656 bp). **c,** Representative Sanger sequencing of *TH*-Cre reporter cell lines showing correct insertion of the Cre cassette in *TH* locus. **d,** Genotyping PCR for 5′ arm (hTH3 UTR 5arm 2F, hTH3 UTR 5arm 1R 1340 bp) and 3′ arm (hTH3 UTR 3arm 2F/PGK hTH3 UTR 3arm 2R 1389 bp) of IRES-Cre cassette inserted into IRES-Cre hPSC clones (B2 and C4). **e,** Genotyping PCR showing homozygous targeted alleles of IRES-Cre cell lines (B2 and C4; knock-in 5640 bp and wild type ~2200 bp). **f,** Representative Sanger sequencing of IRES-Cre reporter cell line showing correct insertion of the IRES-Cre cassette in *TH* locus.

**S2.**
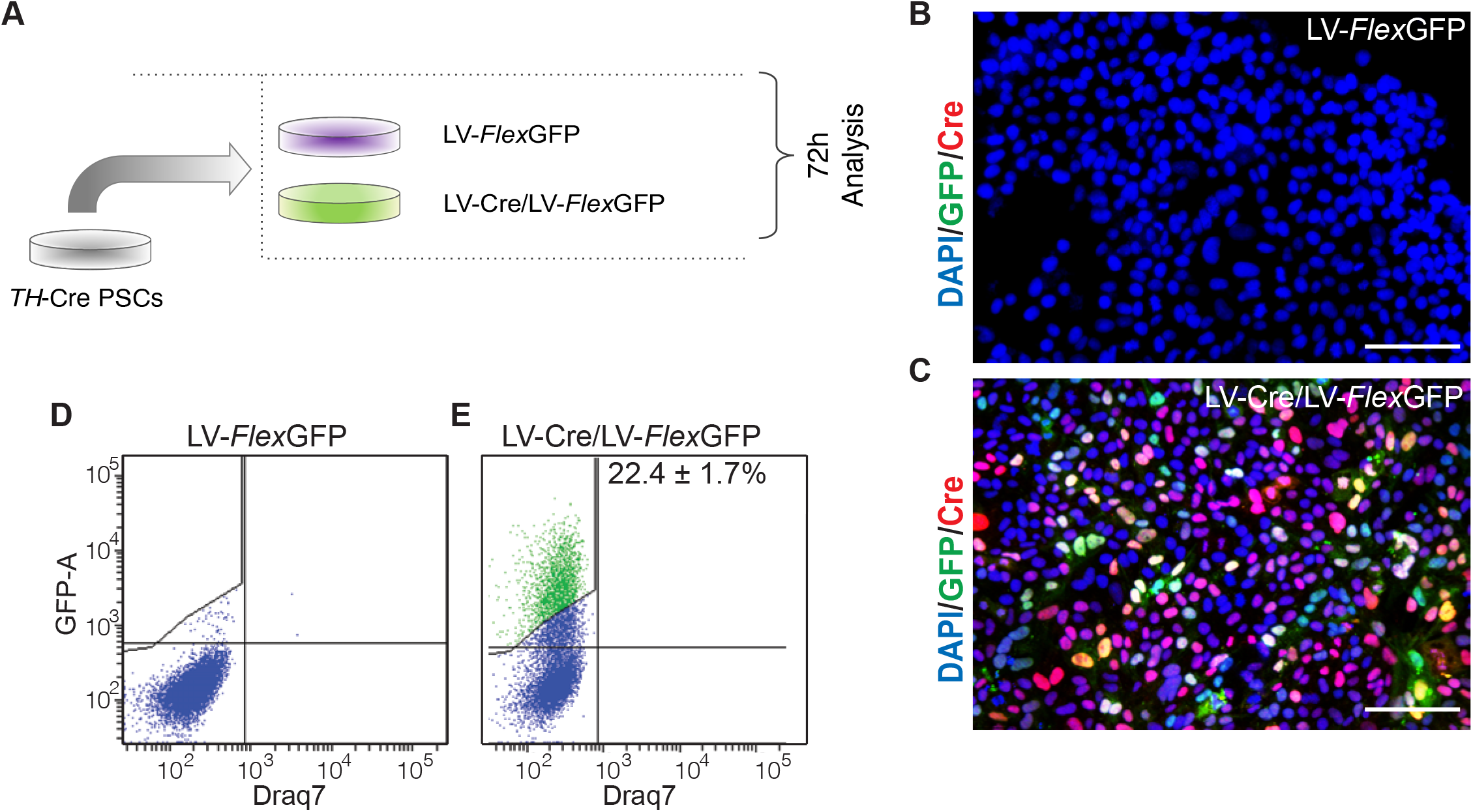
**a,** Schematic overview of experimental plan. **b,** Immunohistochemistry of GFP/Cre in hPSC culture transduced only with LV-FLEx-GFP or **c,** co-transduced with LV-Cre and LV-FLEx-GFP concomitantly. Scale bars, 250 μM. Nuclei were stained with DAPI. **d,** FACS analysis showing GFP+ cell quantification in LV-Cre or LV-FLEx-GFP transduced hPSCs. Data represent mean ± SEM (*n*=3). **e,** Immunohistochemistry of GFP/TH in IRES-Cre hPSC clones (B2 and C4) at day 22 and day 28 during VM differentiation. Scale bars, 250 μM. Nuclei were stained with DAPI.

**Supplementary Table 1.**
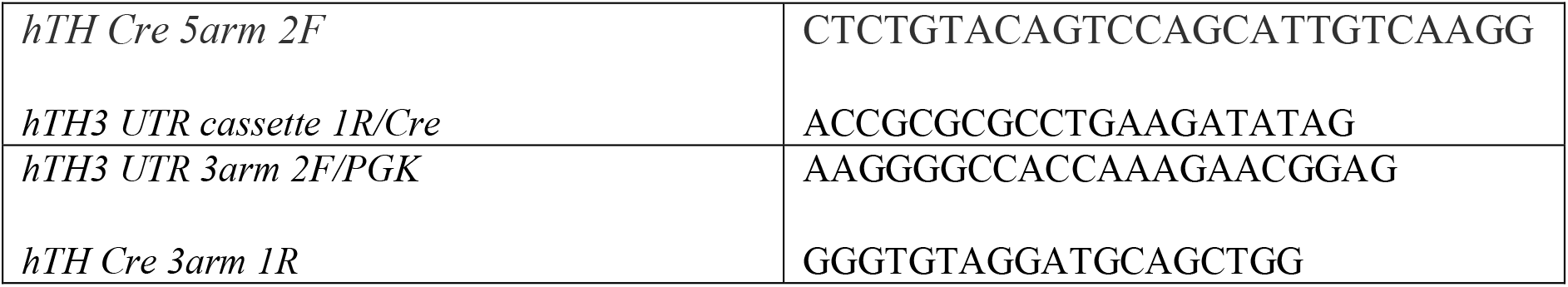
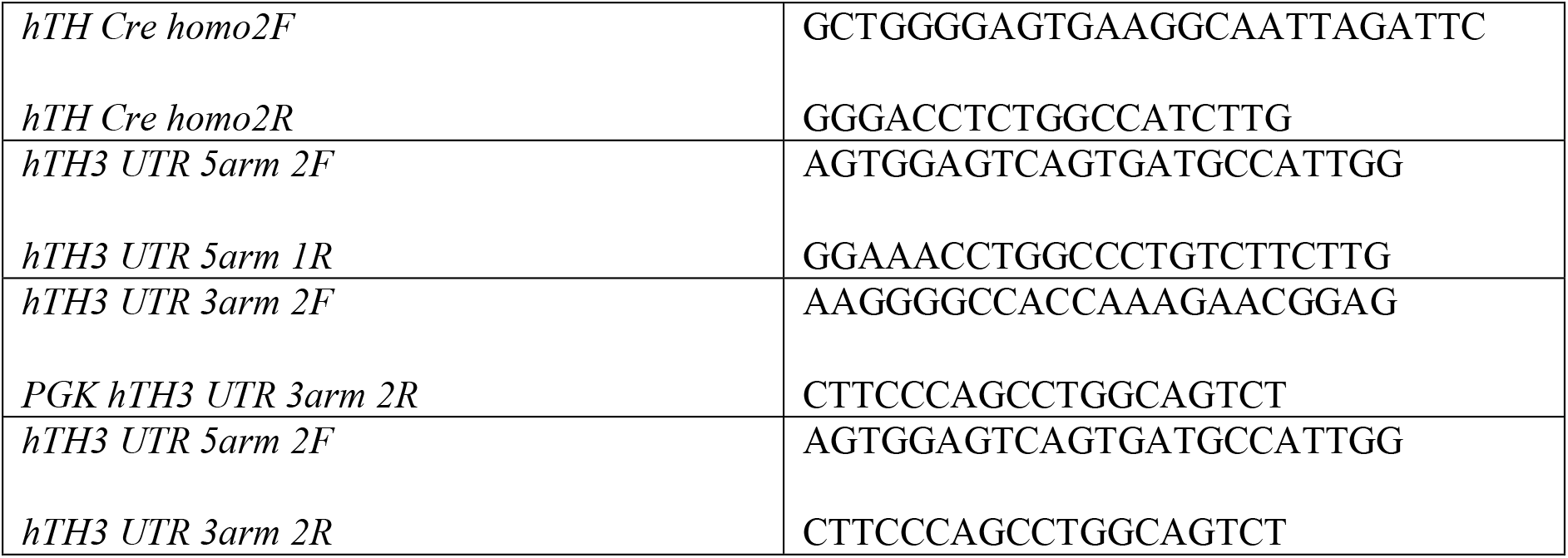

**Supplementary Table 2.**
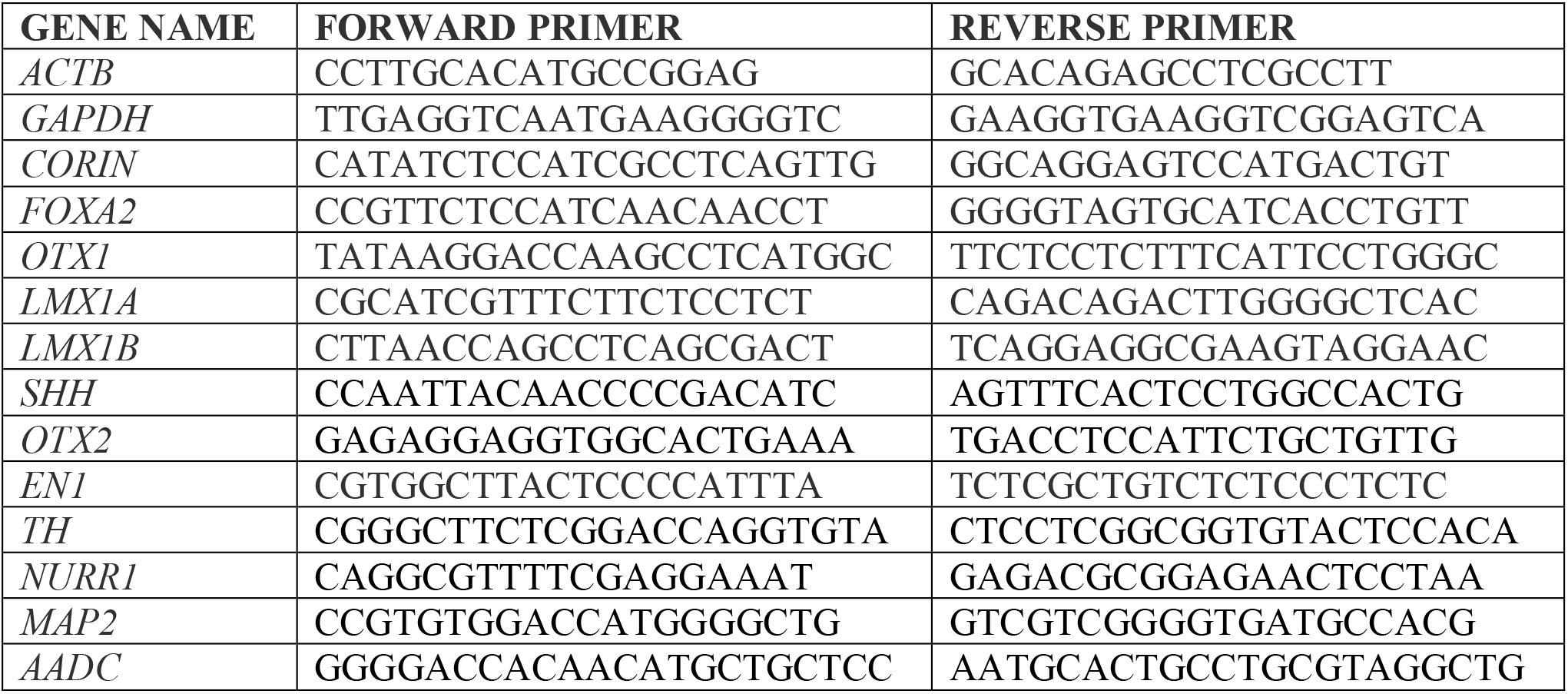
Sequence of qPCR primers

## Methods

### hPSC culture

Undifferentiated RC17 (Roslin Cells, #hPSCreg RCe021-A) and *TH*-Cre hPSC cells were maintained on 0.5 μg/cm^2^ Laminin-521 (Biolamina, #LN-521)-coated plates in iPS Brew medium (Miltenyi, #130-104-368).

### Generation of *TH*-Cre hPSC line

The Cre-recombinase sequence has been inserted immediately after the last codon in exon 1 of TH gene, and therefore gRNAs were designed to target intron 1. Among the gRNA candidates, one of them was selected based on the proximity to the target site and off-target profiles: MS405.TH.g38 (5’-CACATCCACGCCGCGTCCCA**GGG-**3’; in bold is reported Protospacer Adjacent Motif (PAM) site). The selected gRNAs was cloned into a gRNA/Cas9 expression vector by inserting double-stranded oligo cassettes of their DNA sequences between the two Bbs I sites. gRNA constructs were verified by RE digestion and sequencing. After the double-stranded oligos of the two gRNAs were cloned into the gRNA expression vector pBTU6-Cas9-2A-GFP, each construct was transfected into K652 cells individually. Genomic DNA was extracted and PCR was performed (5’-GTGTCTGAGCTGGACGCC-3’; 5’-GATTCAAGATGAAACAAGACACAG-3’). The PCR products were subjected to NGS to assess the frequency of NHEJ events that result from DSB at the HT locus. The cells RC17 cells were transfected with the gRNA MS405.TH.g38 and the donor plasmid. After a 2-week puromycin selection, clones were isolated and genotyped.

### hPSC differentiation

RC17 and *TH*-Cre hPSC cells were differentiated into VM-patterned progenitors using our GMP-grade protocol (Nolbrant et al, Nat Prot 2017). VM-patterned progenitors were either transplanted after 16 days of differentiation or were replated on day 16 as described (Nolbrant et al, Nat Prot 2017) and kept in culture until day 50 of differentiation. For nuclear expression of GFP, *TH*-Cre hPSC cells (passage were transduced with a lentiviral construct under control of the human *TH*-Cre promoter (Addgene, plasmid # …..). The cells were transduced at a multiplicity of infection (MOI) of 1 while cultured on Laminin-521 in iPS-Brew.

### Flow cytometry

*TH*-Cre hPSC and *TH*-Cre VM-patterned single cell suspensions were obtained using Accutase. After dissociation, the cells were analyzed and sorted by FACS based on GFP expression using a BD FACSAria III Cell Sorter (BD Biosciences).

### Animals

Athymic nude female rats (180 g) were purchased from Harlan laboratories (Hsd:RH-Foxn1_rnu_) and housed in individual ventilated cages under a 12:12 hour dark-light cycle with *ad libitum* access to sterilized food and water. All procedures on research animals were performed in accordance with the European Union Directive (2010/63/EU) and approved by the local ethical committees at Lund University and the Swedish Board of Agriculture (Jordbruksverket). For all surgical procedures, rats (minimum 225 g/16–18 weeks) were anesthetized via i.p. injection using a 20:1 mixture of fentanyl-dormitor (Apoteksbolaget). The animals were rendered hemiparkinsonian via unilateral injection of 4 μL of the neurotoxin 6-hydroydopamine at a concentration of 3.5 μg/μL (calculated from free-base HCl) aimed at the following stereotaxic coordinates (in mm): anterior −4.4, medial −1.2, and dorsal −7.8, with the incisor bar set at 2.4. Injections were performed as described (Heuer et al., Exp Neurol, 2013) at an injection speed of 1 μL/minute with an additional 3 min allowed for diffusion before careful retraction of the needle. Intrastriatal transplantation was performed by injecting a total of 300,000 hESC-derived VM-patterned cells resuspanded in 4 μL over four 1 μL deposits (75,000 cells per deposit) at the following coordinates relative to bregma (in mm): anterior +1.2, medial −2.6, dorsal −4.5 and −4.0 and anterior +0.5, medial −3.0. At each deposit the injection cannula was left in place for an additional 2 min before careful retraction to allow for settling of the tissue. Rats were transcardially perfused using 50 mL physiological saline solution (8.9% saline) for 1 minute followed by 250 mL 4% paraformaldehyde (PFA, pH 7.4) solution for 5 min.

### qRT-PCR

Total RNAs were isolated using the RNeasy Micro Kit (QIAGEN, #74004) and reverse transcribed using random hexamer primers and Maxima Reverse Transcriptase (Thermo Fisher, #K1642Invitrogen). cDNA was prepared with SYBR Green Master Mix (Roche, #04887352001) using the Bravo instrument (Agilent) and analyzed by quantitative PCR on a LightCycler 480 supplier using a 2-step protocol with a 60 °C annealing/elongation step. All quantitative reverse transcriptase PCR (qRT-PCR) samples were run in technical triplicates and results are given as fold change over undifferentiated hPSCs. Details and list of primers are reported in Supplementary Table 1.

### Immunocytochemistry

Terminally differentiated cell cultures (day 50) were fixed in 4% PFA for 15 min and then washed three times with PBS. For immunocytochemistry, the cells were blocked for 1–3 hours in PBS + 5% donkey serum + 0.1% Triton X-100 before adding the primary antibodies solution. Primary antibodies were: rabbit anti-TH (1:1000, Merck Millipore, #AB152), anti-mouse anti-Cre (1:500 ab24607), chicken anti-GFP (1:1000, Abcam, #ab13970). After incubation with primary antibodies overnight at 4 °C, the cells were washed three times before adding fluorophore-conjugated secondary antibodies (1:200, Jackson ImmunoResearch Laboratories) and DAPI (1:500). The cultures were incubated with secondary antibodies for 2 hours and finally washed three times.

### Immunohistochemistry

After perfusion, brains were post-fixed for 24 hours in 4% PFA and then cryopreserved in a 30% sucrose solution before being sectioned coronally on a freezing sledge microtome at a thickness of 35 μm in series of 1:8 or 1:12.

Immunohistochemistry was performed on free-floating sections and all washing steps were carried out with 0.1 M phosphate buffered saline with potassium (KPBS, pH 7.4). The sections were washed three times and then incubated in Tris-EDTA pH 8 for 30 min at 80 °C for antigen retrieval. After washing an additional three times, the sections were incubated in blocking solution (5% serum, 0.25% Triton X-100) for 1 hour, before adding the primary antibody solution. Primary antibodies used were: mouse anti-HuNu (1:200, Merck Millipore, #MAB1281), rabbit anti-TH (1:1000, Merck Millipore, #AB152) and chicken anti-GFP (1:1000, Abcam, #ab13970). After incubation with primary antibodies overnight at room temperature, the sections were washed twice and incubated in blocking solution for 30–45 min. For fluorescent immunolabeling, the sections were then incubated with fluorophore-conjugated secondary antibodies (1:200, Jackson ImmunoResearch Laboratories) for 2 hours at room temperature, washed three times and then mounted on gelatin-coated slides and cover-slipped with PVA-DABCO containing DAPI (1:1000).

### Microscopy

Images were captured using either an Epson Perfection V850 PRO flatbed scanner, a Leica DMI6000B widefield microscope, or a Leica TCS SP8 confocal laser scanning microscope. Image acquisition software was Leica LAS X and images were processed using Volocity 6.5.1 software (Quorum Technologies) and Adobe Photoshop. Any adjustments were applied equally across the entire image and without the loss of any information.

### Statistical Analysis

Statistical significance was determined by a two-tailed paired Student’s t-test. P-values <0.05 were considered as statistically significant. Error bars represent SEM.

